# CRISPR-dependent base editing screens identify separation of function mutants of RADX with altered RAD51 regulatory activity

**DOI:** 10.1101/2023.06.19.545603

**Authors:** Madison B. Adolph, Atharv S. Garje, Swati Balakrishnan, Florian Morati, Mauro Modesti, Walter J. Chazin, David Cortez

## Abstract

RAD51 forms nucleoprotein filaments to promote homologous recombination, replication fork reversal, and fork protection. Numerous factors regulate the stability of these filaments and improper regulation leads to genomic instability and ultimately disease including cancer. RADX is a single stranded DNA binding protein that modulates RAD51 filament stability. Here, we utilize a CRISPR-dependent base editing screen to tile mutations across *RADX* to delineate motifs required for RADX function. We identified separation of function mutants of RADX that bind DNA and RAD51 but have a reduced ability to stimulate its ATP hydrolysis activity. Cells expressing these RADX mutants accumulate RAD51 on chromatin, exhibit replication defects, have reduced growth, accumulate DNA damage, and are hypersensitive to DNA damage and replication stress. These results indicate that RADX must bind RAD51 and promote RAD51 ATP turnover to regulate RAD51 and genome stability during DNA replication.

## Introduction

RAD51 is a recombinase that is essential for maintaining genome stability by promoting both double-strand break (DSB) repair and replication fork stability through the formation of a helical nucleoprotein filament on ssDNA [1]. The stability of the RAD51 nucleoprotein filament is highly regulated since it is a key determinant of function. For example, the tumor suppressor BRCA2 stabilizes RAD51 on DNA while helicases remove RAD51 [2-5]. Stable RAD51 filaments are required for strand invasion during homologous recombination (HR); however, these RAD51 filaments must then be disassembled to complete HR [6]. Metastable RAD51 filaments are necessary to promote replication fork reversal at stalled replication forks, a process that involves the annealing of parental DNA strands and the displacement of the nascent strands to form a DNA duplex [7, 8]. The end of the DNA duplex is susceptible to nucleases, but BRCA2-stabilized RAD51 filaments at the reversed fork prevent nascent strand degradation [8-10]. This process is important to maintain genome stability and influences cancer cell responses to drugs that target replication [11].

We previously characterized RADX as a regulator of RAD51 filament stability. RADX is a single stranded DNA (ssDNA) binding protein containing oligosaccharide/oligonucleotide (OB fold) domains [12-14]. RADX binds RAD51 and forms homo-oligomers [12, 15-17]. The second OB fold domain is important for its interaction with ssDNA [12-14], the third is involved in its interaction with RAD51 [15, 18], and a fourth predicted by Alphafold promotes oligomerization [16, 19, 20]. The first predicted OB-fold remains uncharacterized. Each of the characterized RADX biochemical activities is important to regulate replication and replication stress responses.

RADX inactivation leads to slow replication elongation and increased replication fork collapse [12, 13]. Both phenotypes are rescued by inactivating RAD51, suggesting that aberrant RAD51 activity produces these problems. Indeed, RADX decreases RAD51 nucleoprotein filament stability and reduces the amount of RAD51 at replication forks [12, 15, 17, 18]. This function is dependent on its direct interaction with ATP-bound RAD51 on ssDNA leading to a net disassembly of the RAD51 nucleoprotein filament [15]. This regulatory activity allows RADX to inhibit inappropriate RAD51-dependent fork reversal and prevents fork collapse in unstressed cells yet promotes the formation of reversed fork structures in cells experiencing persistent replication stress [18]. While both functions are dependent on RADX binding to RAD51, the mechanisms of how RADX regulates RAD51 to either inhibit or promote replication fork reversal depending on cellular context remain unclear.

CRISPR-based genetic screening methods have been widely used to study loss of function, gain of function, and separation of function phenotypes [21]. While CRISPR-screens initially utilized Cas9 and a guide RNA (sgRNA) to generate DNA double-strand breaks (DSBs) in a sequence specific manner to disrupt gene function, newer CRISPR-based methods that are DSB independent have been developed that generate targeted mutations at specific DNA bases. For example, one approach employs the fusion of a mutant Cas9 (Cas9 D10A) with the rat cytosine deaminase rAPOBEC1 (BE3) to introduce C to T transitions within a defined base editing window [22, 23]. This base editing method has been applied to generate mutations across genes such as *BRCA1* and *MAP2K1*, and recently was used to assess nucleotide variants in a high-throughput manner across 86 DNA damage response genes [24-27].

Here, we utilized this CRISPR-dependent base editing approach with a library of sgRNAs targeted across *RADX* to delineate which of its domains and activities are needed to regulate DNA replication and replication stress responses. We identified a cluster of separation of function mutations on an interdomain helix of RADX in the Alphafold model that selectively reduces its ability to stimulate RAD51 ATP-hydrolysis while retaining the ability to oligomerize, bind ssDNA and RAD51, and localize to replication forks. Cells expressing these mutants accumulate RAD51 on chromatin, accumulate DNA damage, have reduced viability and are hypersensitive to replication stress. These results indicate that stimulating RAD51 ATPase activity is an important RADX function during DNA replication.

## Results

### CRISPR dependent base editing identifies regions of interest in RADX

We were interested in characterizing RADX to better understand how it functions to regulate replication and genome stability. Many genetic screens to study replication and repair factors utilize replication stress inducing or DNA damaging agents since inactivating these factors often causes hypersensitivity to these agents [25, 28, 29]. However, even complete inactivation of RADX does not cause marked hypersensitivity to these agents [12]. Therefore, we developed two alternative RADX loss of function screening strategies.

First, since RADX inactivation causes an increase in DNA damage during S-phase, we hypothesized that it may cause a cell growth disadvantage in p53-proficient RPE1 cells immortalized with hTERT (hereafter referred to as RPE WT cells). To test this hypothesis, we used a two-color growth competition assay in RPE WT cells expressing GFP or mCherry transfected with siRNAs targeting RADX (siRADX) or non-targeting controls (siNT), respectively. The ratio of mCherry to GFP cells was monitored over time. As predicted, cells transfected with RADX siRNA were depleted from the population, leading to a 6-fold increase in siNT expressing cells (Figure 1A). In contrast, RADX inactivation does not impair the growth of RPE p53^-/-^ cells that lack the p53 cell cycle checkpoint (Figure 1A).

**Figure 1.**
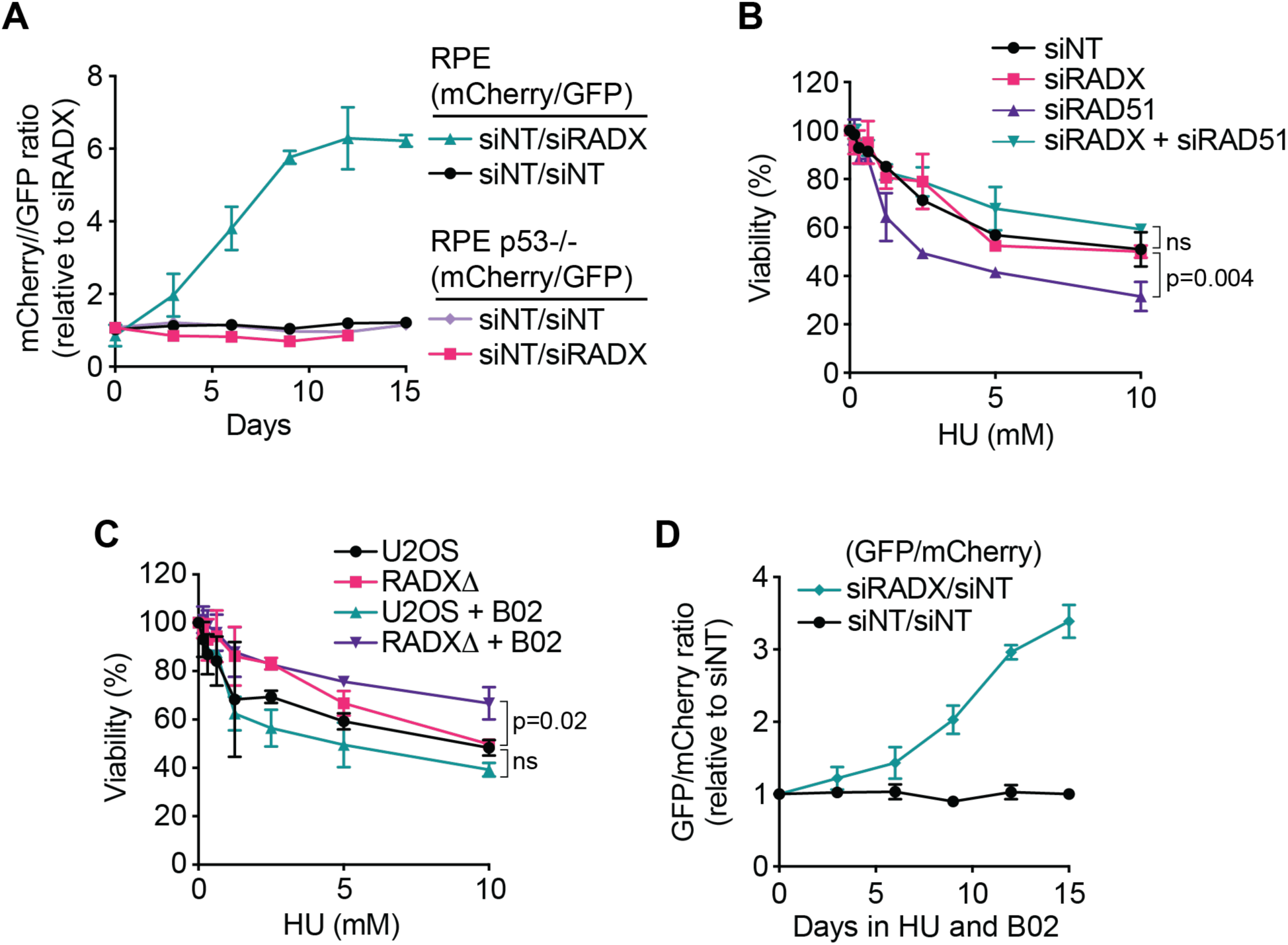
RADX inactivation decreases RPE WT cell growth but suppresses RPE p53^-/-^ cell sensitivity to RAD51 inhibition in the presence of replication stress. **A)** Competitive growth assay in untreated RPE WT and RPE p53^-/-^ cells stably expressing either GFP or mCherry. Data represent cells transfected with non-targeting or siRADX in mCherry or GFP cell lines, respectively, and normalized to the day 1 time point. Each condition was plated in triplicate wells. Mean +/- SD, n=2. **B)** Cells transfected with siRNAs were treated with HU and viability was measured 72 hours later (mean ± SEM, n = 3). p value was calculated using a one-way ANOVA **C)** Parental or RADXΔ cells were treated with 25 mM of B02 over a range of HU concentrations and viability was measured 72 hours later (mean ± SEM, n = 3). p value was calculated using a one-way ANOVA. **D)** Competitive growth assay in RPE p53^-/-^ cells stably expressing either GFP or mCherry and treated with 2.5 mM HU and 25 mM B02. Data represent cells transfected with non-targeting or siRADX in mCherry or GFP cell lines, respectively, and normalized to the day 1 time point. Mean +/- SD for n=2.

As a second screen to measure RADX function, we utilized the observation that RADX inactivation suppresses hypersensitivity of cells with reduced RAD51 to replication stress agents due to a restoration of fork protection [17]. Knocking down RAD51 with siRNAs causes hydroxyurea (HU) hypersensitivity (Figure 1B). However, inactivating RADX at the same time suppresses this sensitivity (Figure 1B). The same effect is observed in RADX knockout cells treated with the RAD51 inhibitor B02 (Figure 1C) [30]. We confirmed these results using the two-color growth competition assay in RPE p53^-/-^ cells which showed 4-fold enrichment of RADX-deficient cells in the presence of HU and B02 compared to controls (Figure 1D). Thus, this B02/HU-treated RPE p53^-/-^ cellular assay generates the opposite growth effect when RADX is inactivated compared to the untreated, wild-type RPE cell assay.

Next, we employed both RADX functional assays in CRISPR-dependent base editing screens. We generated a lentiviral guide RNA (sgRNA) library to tile mutations across the entire *RADX* gene. Additionally, previously published cell lethality controls, known as iSTOP controls, negative controls targeting the *AAVS1* gene and empty window controls were included in the sgRNA library [31, 32]. The sgRNA library (1912 sgRNAs) was transduced into wild-type RPE or RPE p53^-/-^ cells expressing the BE3 base editor. We verified the expression of the BE3 cassette by immunoblotting and through the activity of the uracil DNA glycosylase inhibitor (UGI) present on the cassette (Supplemental Figure 1A-B). Transduced cells were then cultured for 18 days and the representation of the sgRNA library at day 18 (T18) compared to day 0 (T0) was examined by next-generation sequencing (NGS) to quantify the amount of each sgRNA in the populations and generate a log2-fold change (LFC) and p-value. A schematic overview of the screening strategy is presented in Figure 2A. As expected, the positive iSTOP controls were depleted by T18, whereas the *AAVS1* and empty window controls remained unchanged from T0 (Figure 2B). We then examined all sgRNAs to RADX and considered sgRNAs that caused LFC of >+/-0.5 and a p value <0.01 as significant. As expected, many RADX sgRNAs show a marked depletion at T18 in wild-type RPE cells and a corresponding enrichment of these same sgRNAs in RPE p53-/- cells treated with HU and B02 (Figure 2C).

**Figure 2.**
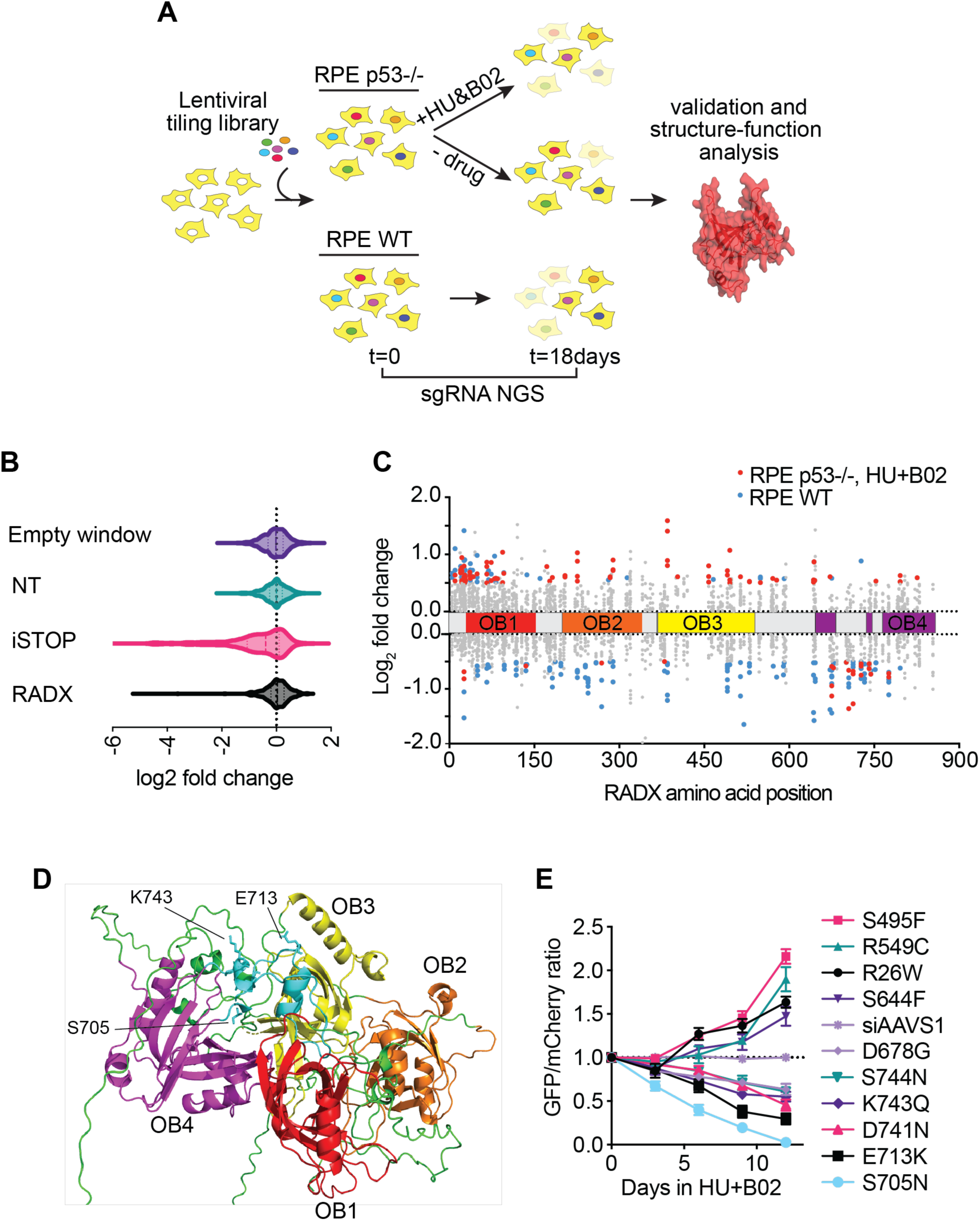
CRISPR-dependent base editing screens identify mutants of interest in RADX. **A)** Schematic of CRISPR-dependent base editing screen. RPE WT or RPE p53^-/-^ cells were transduced with lentiviral sgRNA library and then cultured for 18 days after selection. sgRNA abundance between day 0 (T0) and day 18 (T18) was determined by next-generation sequencing (NGS). **B)** Density plots of LFC for control and RADX gRNAs in RPE WT cells. **C)** Lollipop plot of RADX sgRNAs and their LFC values in untreated RPE WT or RPE p53^-/-^ in the presence of 2.5 mM HU and 10 μM B02. sgRNAs that did not meet LFC or p-value significance cutoffs are colored grey while sgRNAs with LFC >0.5 and p-values >0.01 are colored blue (RPE WT, untreated) or red (RPE p53^-/-^ HU and B02). **D)** Predicted location of RADX 3,4-ID mutants based on Alphafold database structure is colored in teal. Stick model of amino acid residues tested in this region predict they are surface exposed. **E)** Competitive growth assay in BE3-expressing RPE p53^-/-^ cells treated with HU and B02 expressing the indicated sgRNAs. Data represent the GFP-sgRNA mCherry-*AAVS1* control ratio normalized to the day 1 (T1) time point. Mean ± SD for n = 2.

While targeting most RADX domains yielded opposite effects in the two screens as expected, there were two regions of interest that demonstrated unexpected phenotypes. Some sgRNAs targeted at the RADX N-terminus appeared to be enriched in both screens, indicating a growth advantage for these cells. Secondly, sgRNAs targeted between residues 678-750 were depleted across both screens in RPE WT and RPE p53^-/-^ cells, indicating that mutating *RADX* in this region was deleterious for cell growth even in the presence of B02 and HU. In this study we chose to focus on the region of interest from residues 678-740 due to the defects on cell growth they presented. Alphafold predicts these mutations are on a surface between the third and fourth oligosaccharide/oligonucleotide binding domains (OB); therefore, we refer to mutations in this region as OB3-4 interdomain mutants, abbreviated 3,4-ID (Figure 2D). This region has not previously been characterized as important for RADX function.

We confirmed this striking dropout phenotype in RPE p53^-/-^ cells in the presence of HU and B02 by growth competition assays comparing the relevant individual RADX GFP-sgRNAs to an mCherry-*AAVS1* targeting control (Figure 2E). As in the library screen, the individual sgRNAs to the 3,4-ID region reduced cellular fitness in these cells; whereas the sgRNAs targeting other RADX regions were enriched as they confer resistance to HU and B02. We also confirmed that these sgRNAs yielded the predicted mutational outcome by sequencing the *RADX* genomic DNA in these cells (Supplemental Figure 1C).

To determine if any of these mutations were predicted to be deleterious, we examined the variants in this region using predictive tools to determine their functional relevance. Examination of these mutations using MutPred did not predict disruption of protein folding or structural integrity. However, mutations in this region are predicted as likely deleterious by Ensembl variant effect predictor (VEP), Polymorphism Phenotyping v2 (PolyPhen-2) and Sorting Intolerant From Tolerant (SIFT). Notably, variants S705N and E713K were predicted to be the most disruptive within the cluster of variants. Analysis of RADX in cBioPortal shows that *RADX* mutations within this region appear in various cancers presented in the TCGA database, such as Uterine Corpus Endometrial Carcinoma and Cervical Squamous Cell Carcinoma. Most notably E713K, D741N and K743Q appear as variants in at least one TCGA study. Therefore, we prioritized these mutants for additional analyses.

### Cells expressing the 3,4-ID mutants of RADX have slow replication elongation, accumulate DNA damage, and are hypersensitive to DNA damaging agents

We next asked whether these RADX mutations generated any additional phenotypes in cell-based assays in U2OS cells where many RADX functions had previously been characterized. RADXΔ cells derived through CRISPR-Cas9 editing were complemented with lentivirus encoding GFP-tagged wild-type RADX or the various RADX 3,4-ID mutants. We found that the 3,4-ID mutants of RADX have elevated levels of the DNA damage marker γH2AX in S-phase cells similar to RADXΔ cells and in contrast to cells complemented with wild-type RADX (Figure 3A). Since inactivation of RADX leads to slow replication fork elongation we examined fork elongation rates in RADXΔ cells and cells complemented with wild-type or 3,4-ID mutants of RADX. The 3,4-ID mutants of RADX have slow replication fork rates comparable to RADXΔ cells (Figure 3B). The elevated γH2AX and slow replication is not due to lack of expression of the RADX mutants as they all expressed to similar levels as wild-type RADX, and all are overexpressed compared to endogenous RADX (Figure 3C). Similarly, each mutant retains localization to replication forks at comparable levels to wild-type RADX (Figure 3D).

**Figure 3.**
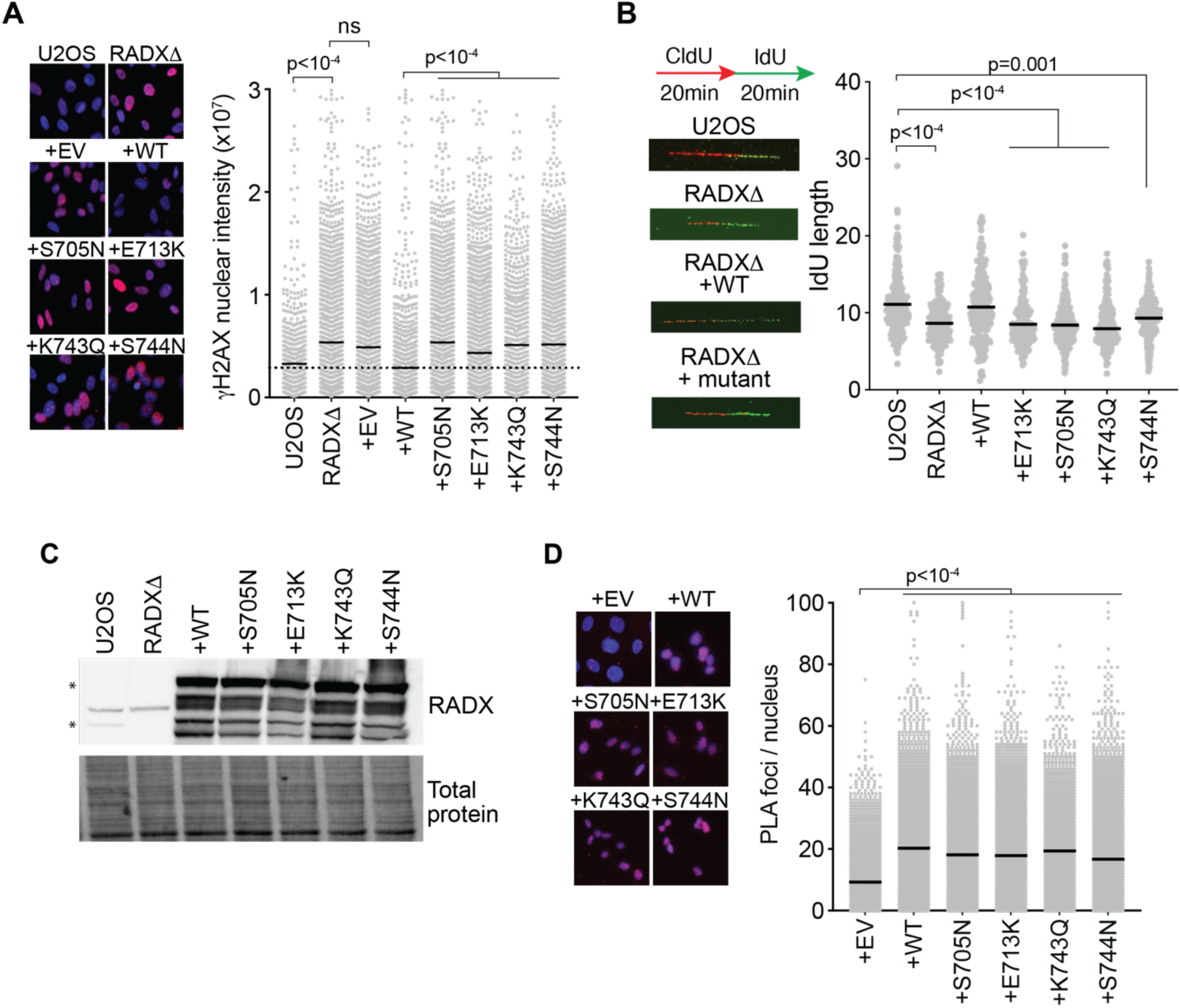
RADX 3,4-ID mutants slow replication elongation and increase DNA damage in S-phase cells. **A)** γH2AX intensity in EdU-positive S phase U2OS wild-type or RADXτι cells complemented with empty vector (EV) control, wild-type RADX, or the indicated RADX mutants. P values were derived from a one-way ANOVA with a Dunnett’s multi-comparison test. **B)** Cells were labeled with CldU and IdU as indicated and then analyzed by DNA combing to measure fork speed. A representative experiment is shown with p values derived from a one-way ANOVA with Dunnett’s multiple-comparison test**. C)** Immunoblots of U2OS or RADXτι cells infected with lentivirus expressing the wild-type (WT) RADX or RADX mutants. **D)** Proximity ligation assay between FLAG-RADX and EdU to measure localization to replication forks. Cells were labeled for 10 min with EdU. P values were derived from a one-way ANOVA with a Dunnett’s multi-comparison test.

Similar to the results in RPE cells, expression of the 3,4-ID mutants of RADX in RADXΔ U2OS cells led to growth of fewer colonies than wild-type RADX expressing cells (Figure 4A). However, RADXΔ U2OS cells do not exhibit this viability defect in U2OS cells (Figure 4A). RADXΔ cells are also not hypersensitive to DNA damaging agents; however, cells expressing 3,4-ID RADX mutants became hypersensitive to multiple DNA damaging and replication stress agents including HU, cisplatin and camptothecin (CPT) (Figures 4B-4D). Thus, expression of the 3,4-ID mutants of RADX is worse for U2OS cell growth and replication stress survival than deletion of RADX. These observations suggest the presence of these mutants that retain localization to replication forks may interfere with alternative compensatory mechanisms that act when RADX is deleted.

**Figure 4.**
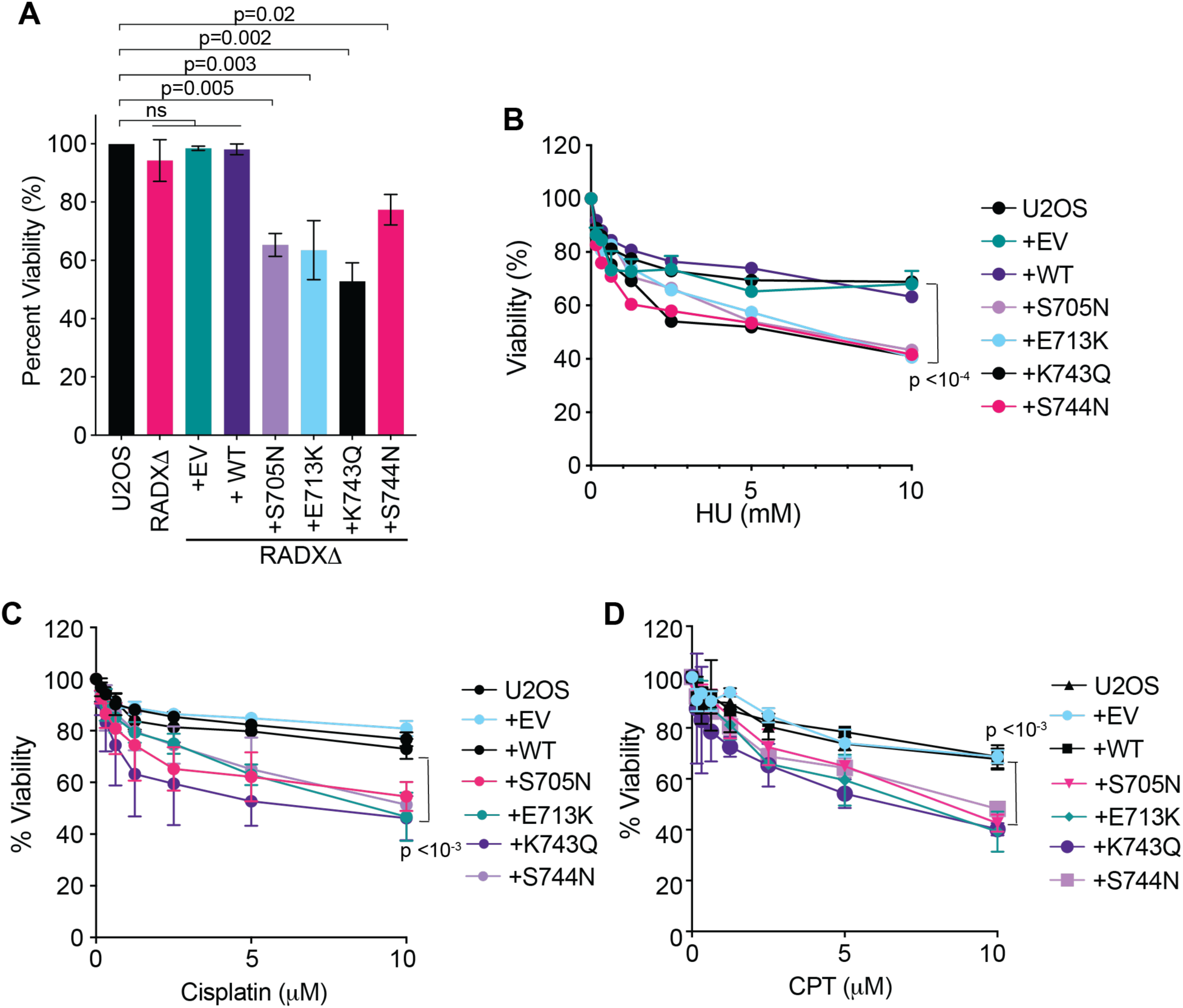
RADX 3,4-ID mutants decrease cell viability and cause hypersensitivity to replication stress and DNA damaging agents. **A)** RADXΔ U2OS cells complemented with empty vector (EV) control, wild-type RADX or the indicated mutants of RADX were plated in the absence of drug and surviving colonies examined by methylene blue staining and quantified (mean ± SEM, n = 3). **B-D)** Parental or RADXΔ cells complemented with empty vector control, wild-type RADX or the indicated mutants of RADX were treated with the indicated drugs and viability was measured 72 hours later (mean ± SEM, n = 3). P values were generated with a one-way ANOVA with a Dunnett’s multi-comparison test.

In unstressed cells, the ability of RADX to prevent replication fork instability, and maintain replication fork elongation is dependent on its interactions with ssDNA, RAD51, and RADX oligomerization [12-18]. However, the residues within the 3,4-ID region are not within the regions previously characterized as necessary for these biochemical activities. Indeed, purified 3,4 ID RADX mutant proteins interact with RAD51 in the presence of ATP comparable to wild-type RADX (Figure 5A). These mutants also interact with ssDNA as expected since the mutations are not in the characterized ssDNA binding domain (Figure 5A). Similarly, epitope-tagged 3,4-ID mutants could co-immunoprecipitate with wild-type RADX indicating they retain the ability to form oligomers (Figure 5B). Therefore, the cellular phenotypes caused by the 3,4-ID RADX mutants do not appear to be attributable to loss of these characterized interactions.

**Figure 5.**
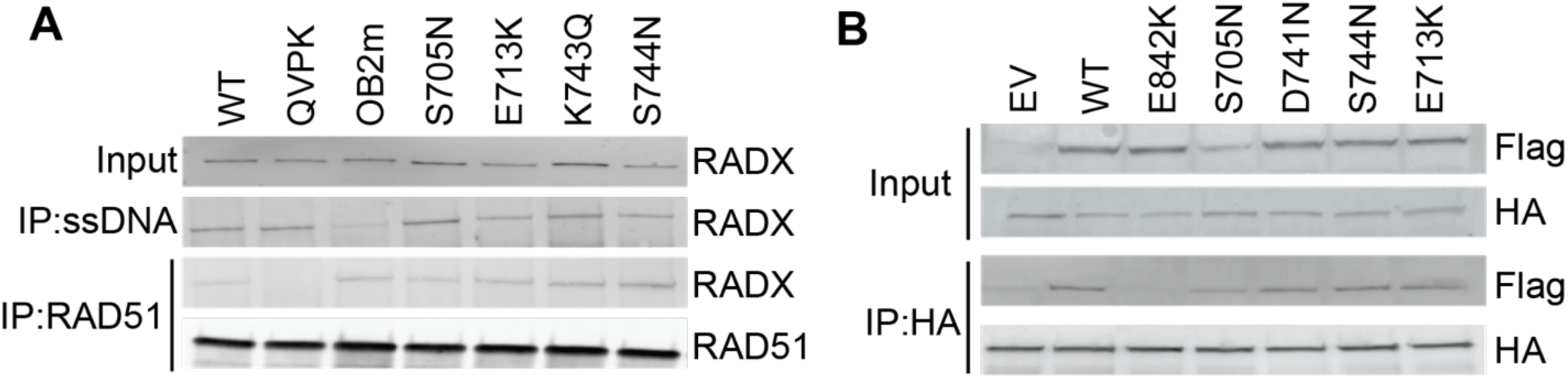
RADX 3,4-ID mutants bind ssDNA, bind RAD51-ATP, and oligomerize. **A)** Interaction of purified RADX and RADX mutants with RAD51 in the presence of ATP, or biotinylated ssDNA. **B)** Flag-RADX wild-type was co-expressed with HA-RADX mutants into HEK293T cells. HA immunoprecipitation from cell lysates was followed by SDS-PAGE and immunoblotting. (EV = empty vector).

### 3,4-ID RADX mutants reduce RAD51 ATP turnover leading to RAD51 accumulation on chromatin

Loss of RADX leads to an increase in RAD51 localization to chromatin in S-phase cells [12]. This increased accumulation of RAD51 results in slower fork speeds which can be rescued by depleting RAD51 [12]. Likewise, the slow replication fork speeds observed in cells expressing the 3,4-ID RADX mutants could be rescued by depletion of RAD51 (Figure 6A). RAD51 levels and number of foci are also increased on chromatin in RADXΛ1 cells complemented with 3,4-ID RADX mutants similar to what is observed in the empty vector expressing cells and in contrast to the reduction of RAD51 observed with wild-type RADX complementation (Figures 6B and 6C). These results suggest that the defects observed for these mutants is tied to some aspect of RAD51 regulation even though the mutants retain the ability to interact with RAD51.

**Figure 6.**
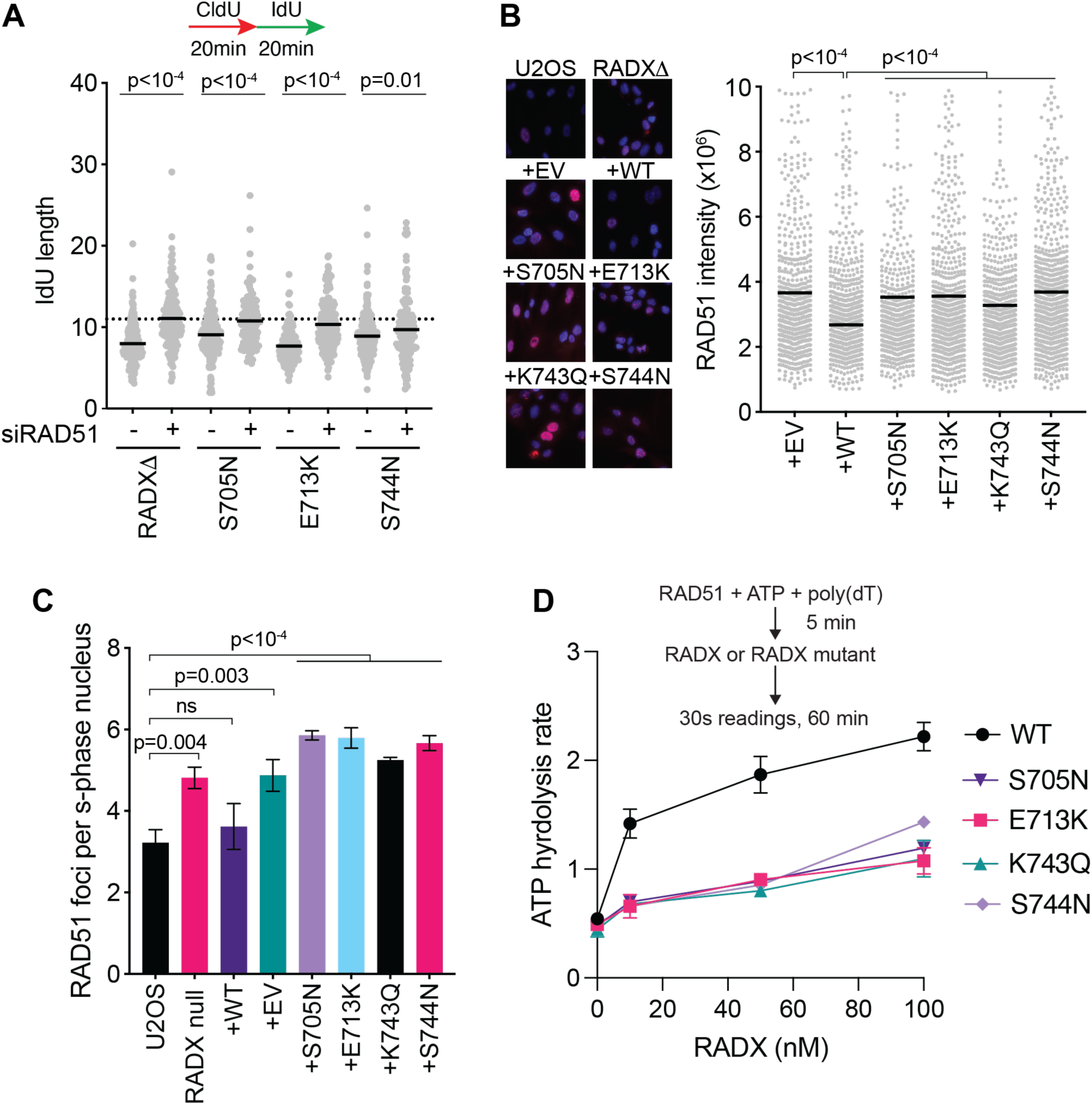
3,4-ID mutants of RADX fail to regulate RAD51. **A)** Parental or RADXΔ cells complemented with empty vector control, wild-type RADX or the indicated mutants of RADX were transfected with siNT or siRAD51 and labeled with CldU and IdU as indicated and then analyzed by DNA combing for replication fork speed. Representative experiment is shown with p values derived from a one-way ANOVA with Dunnett’s multiple-comparison test. **B and C)** RADXΔ U2OS cells complemented with empty vector control, wild-type RADX or the indicated mutants of RADX were labeled with 10 μM EdU for 30 min. Cells were stained for RAD51 and EdU after detergent extraction. **B)** The intensity of chromatin-bound RAD51 in EdU-positive cells is quantified. **C)** The number of foci per S-phase nucleus for RAD51 is quantified. p values were derived from a Mann-Whitney test. **D)** RAD51 ATPase assay with increasing concentrations of RADX added to the reaction after the addition of RAD51.

To further understand why the 3,4-ID mutants lead to RAD51 accumulation, we examined the ability of wild-type and mutant RADX to stimulate RAD51 ATP hydrolysis. RAD51 ATP hydrolysis promotes its removal from DNA [33]. Using purified proteins, we observed that the RADX 3,4-ID mutants were unable to stimulate the RAD51 ATP hydrolysis rate to similar levels as wild-type RADX (Figure 6D) consistent with the increased RAD51 retention on DNA in S-phase cells expressing these mutants. Thus, the cellular phenotypes of these 3,4-ID mutants, including accumulation of toxic RAD51 at forks, correlate with their inability to stimulate RAD51 ATP hydrolysis.

## Discussion

This work highlights the ability of CRISPR-dependent base editing screens to identify separation of function mutants in proteins of interest. Using complementary screening assays, we identified a cluster of mutations on a surface of RADX that separates the ability of RADX to bind RAD51, bind ssDNA, and oligomerize from its ability to stimulate RAD51 ATP hydrolysis. Expression of these mutants leads to decreased clonogenic survival and hypersensitivity to DNA damaging agents. We conclude that an important function of RADX is to stimulate RAD51 ATP hydrolysis which helps to remove RAD51 from DNA. Interestingly, some phenotypes caused by expression of these mutants including the hypersensitivity to replication stress are not observed with complete loss of RADX expression. Since these mutants retain the ability to bind DNA, bind RAD51, and localize to replication forks, it is likely that they prevent alternative mechanisms from acting as might happen in cells lacking RADX altogether.

Exactly how RADX stimulates RAD51 ATPase activity remains unclear. It may induce an alteration in the active site of RAD51 in a way that promotes ATP turnover, similar to other RAD51 modulators like Srs2 [15, 34-36]. The 3,4-ID mutants may bind RAD51 differently than wild-type RADX, or with different affinity leading to differences in ATP turnover. Alternatively, these RADX mutants may bind the RAD51 nucleoprotein filament in a manner that traps RAD51 on ssDNA, preventing the intrinsic dynamics of RAD51 [37] and partially mimicking other circumstances in which RAD51 forms a hyperstable filament. For example, the RAD51 K133R mutation that largely abrogates its ATP hydrolysis activity generates hyperstable filaments that cause HR deficiency and hypersensitivity to DNA damaging agents [33, 38]. There are numerous other modulators of RAD51 that control rates of ATP hydrolysis. BRCA2 decreases ATP hydrolysis to stabilize the nucleofilament, whereas FIGNL1 and PARI lead to increased ATP hydrolysis and net dissociation of the RAD51 filament [5, 39, 40]. The conformational change between ATP bound RAD51 and ADP bound RAD51 transitions the nucleofilament from an active to inactive state, respectively. These conformational changes are required for RAD51 to fulfill its functions in repair and replication.

Our results indicate that stimulation of RAD51 ATP hydrolysis by RADX is essential for regulating RAD51 and maintaining genome integrity. RADX is expressed at much lower levels than RAD51 and competes with BRCA2 for RAD51 binding [15]. An attractive model is that RADX caps the end of a RAD51-ssDNA filament and promotes the dissociation of RAD51 from ssDNA by promoting conversion of RAD51 filaments to the inactive ADP-bound form. A high-resolution structure of RADX in complex with a RAD51 nucleofilament will provide critical insight to the nature of this interaction and the molecular mechanism by which RADX functions.

In summary, we identified separation of function mutants of RADX using a CRISPR-dependent base editing screen. Characterization of these mutants showed that the ability of RADX to stimulate ATP hydrolysis by RAD51 is important to control DNA replication and replication stress responses. Interestingly, several of these mutations are found in human tumors where they could not only drive genome instability but also make those tumors hypersensitive to some DNA damaging chemotherapies.

## Acknowledgements

This research was supported by NIH grants R01GM116616 to D.C., R35GM118089 to W.J.C., and P01CA092584 to D.C. and W.J.C. RADX proteins were generated with the assistance of the SBDR EMB core (P01CA092584). M.M. is supported by the French Ligue against Cancer (équipe labellisée).

## Author contributions

M.B.A. and D.C. conceived of the project. A.S.G helped with molecular biology and western blot experiments. S.B and W.J.C contributed to experimental design. M.B.A. completed all other experiments, which D.C. supervised. F.M. and M.M. provided RAD51 protein. M.B.A. and D.C. wrote the manuscript with input from the other authors.

## Materials and Methods

### Cell culture

U2OS, and HEK293T cells were cultured in DMEM with 7.5% fetal bovine serum (FBS). hTERT-RPE1 cells were cultured in DMEM:F12 supplemented with 7.5% FBS. hTERT-RPE p53 -/- cells were a gift from Daniel Durocher [29] and were cultured in DMEM with 7.5% FBS, 1X Glutamax and 1X non-essential amino acids. Cells were cultured at 37°C and 5% CO_2_ with humidity. All cell lines were regularly tested for mycoplasma and verified using short tandem repeat profiling.

### Cell line generation

U2OS RADXτι were described previously [12]. Complementation of RADXτι cells with cDNA expression vectors of mutants of interest was completed by lentiviral infection and selection for the linked puromycin resistance cassette as described [12]. Plasmid and siRNA transfections were performed with polyethylenimine for plasmid overexpression in 293T and Dharmafect1 (Dharmacon) for siRNA in all other cell lines.

### Generation of BE3 expression cell lines and validation

Cells expressing the BE3 cassette (BE3-FNLS-P2A-BlastR) [31, 41] were generated as described in [25]. Briefly, cells expressing the BE3 cassette were generated by lentiviral infection of RPE WT and RPE p53^-/-^ and BE3 expressing cells were selected in blasticidin at 10 μg/mL for 72 hours. BE3 cell lines were then seeded at low density for single cell clone recovery. Flag-BE3 expression was determined by immunoblot. BE3 expressing cells were further validated for UGI activity by harvesting and lysing cells in 50 mM Tris-Cl pH 7.4, 1% Igepal, 0.1% sodium deoxycholate, 10% glycerol, 150 mM NaCl, 1 mM DTT, 20 μg/ml RNaseA with an EDTA-free protease inhibitor cocktail tablet (Roche). Cell lysates (or buffer) were incubated with 100 nM of a 42-nucleotide ssDNA oligonucleotide with a single deoxyuridine and a 5′ fluorescein tag (5′ (6-FAM)-CTA TGA TGA CTC TTC TGG TCU GGA TGG TAG TTA AGT GTT GAG 3′) at 37°C for 30 minutes. As a control, UDG from *E. coli* (New England Biolabs) was added to one reaction for complete uracil removal. DNA was then heated at 95°C for 10 min in the presence of 0.2 M NaOH to cleave abasic sites. Samples were mixed with an equal volume of formamide/EDTA loading buffer, resolved on a 10% TBE-Urea gel, and scanned using a Typhoon imager (GE Healthcare).

### Plasmids

The RADX cDNA was obtained from the ThermoScientific Open Biosystems Human ORFeome collection (Catalog number OHS-1770). The mutant RADX OB2m and RADX QVPK were described previously [12, 15, 17]. BE3-FNLS-P2A-BlastR was generously provided by Alberto Ciccia and is previously described [31]. pLenti-Guide-Puro (Addgene #52963, [42], was utilized for sgRNA expression. For characterization of individual sgRNAs, selected RADX sgRNAs were cloned into a modified pLenti-Guide-Puro vector where the Cas9 was replaced with either GFP-NLS or mCherry-NLS (LentiGuide-puro-GFP-NLS and LentiGuide-puro-mCherry-NLS) as previously described [42]. A sgRNA targeting the *AAVS1* gene was used as a control.

### Antibodies and siRNAs

Antibodies used in this study include Flag (Sigma F7425), HA (Biolegend, 901501), GFP (Abcam, ab13970), RAD51 (14B4, Abcam), RADX (NBP2, Novus), γH2AX (JBW301, Millipore), IdU (Abcam Cat#ab6326), and CldU (BD Cat#347580). siRNAs include ON-TARGETplus CXorf57 siRNA (Dharmacon J-014634-21) and RAD51 siRNA (Dharmacon J-003530-12).

### sgRNA library design and generation

A custom DNA library was designed to tile mutations across the entire *RADX* gene. All possible 20bp sequences upstream of NGG PAM sites and within the base editing window (13-18 nucleotides from the PAM) were designed using Benchling using the *hg38* human genome assembly. Negative controls included sgRNAs targeting the *AAVS1* locus, non-targeting controls from the GeCKO_v2 library and “empty window” controls consisting of sgRNAs targeting common essential genes that lack editable bases in the BE3 base editing window [25]. Positive controls included iSTOP cell lethality controls as described [43]. Control sgRNAs were designed to represent 10% percent of the final library. Duplicated sgRNAs or sgRNAs containing the BsmBI site used for cloning were removed from the library. Oligonucleotides were designed including a BsmBI restriction site for cloning into pLentiGuide-Puro and primer sites were added for specific library amplification within the pool. The final sequence of the sgRNA design was: TTTCTTGGCTTTATATATCTTGTGGAAAGGACGAAACACCG(sgRNA)GTTTTAGAGCTAGAAATAGCAAGTTAAAATAAGGCTAGTCCGT. The sgRNA library was generated by GenScript and QC was done by next generation sequencing (NGS) to ensure equal representation of the sgRNAs in the library.

### BE3 base editing screen

Both screens were performed in biological duplicates in both a pooled BE3 cell line and two single cell clones totaling n=6 for each screen. The general protocol for the screen was adapted from [25, 29]. Library coverage of at least 1000 cells per sgRNA was preserved at every step. RPE WT and RPE p53^-/-^ were transduced with the custom lentiviral library at a low MOI (0.3). Media was changed after 24 hours. After 48 hours media containing puromycin was added to a concentration of 20 mg/mL and maintained for 48 hours. For the duration of the screen cells were maintained in selection media containing 15 mg/mL puromycin and 5 mg/mL blasticidin. After 48 hours of selection, cells were harvested for T0 sgRNA representation and replated into conditions either with or without the addition of drug as described in Figure 2A. Drug was refreshed every 3 days, and cells were collected at T18 for final sgRNA library representation. Hydroxyurea (HU) was used at a concentration of 2.5 mM and the RAD51 inhibitor B02 was used at a concentration of 10 μM. Drug concentrations were optimized to achieve a lethal dose 20 (LD20) assaying cell viability every 3 days for 18 days. Genomic DNA was isolated from cells harvested at T0 and T18 using the QIAamp Blood Midi kit (Qiagen). Integrated sgRNA sequences were PCR amplified with Q5 high-fidelity DNA polymerase (New England Biolabs). Briefly, 6 μg of genomic DNA was amplified per condition using a staggered forward primer mix and a unique reverse primer as described in [25]. The PCR amplification was as follows: 30 s at 98°C; followed by 10 s at 98°C, 30 s at 58°C, 20 s at 72°C for 20 cycles; and 2 min at 72°C. Samples were then treated with ExoI to remove any contaminating primers and PCR clean-up was performed with the Qiagen PCR clean up kit. Samples were submitted to the Vanderbilt Technologies for Advanced Genomics (VANTAGE) Next Generation Sequencing (NGS) core. Samples were barcoded with Illumina dual indexes (UDI) and the products were separated on 2% agarose gels, gel purified, multiplexed and sequenced on a Novaseq sequencing platform.

### Two color competition assays

U2OS cells were infected with lentiviral LentiGuide-puro-mCherry-NLS or LentiGuide-puro-GFP-NLS to generate stable cell lines expressing mCherry or GFP. These cells were then transfected with siNT or siRADX to generate GFP-siRADX and mCherry-siNT cells. 48 hours after transfection cells were plated for the experiment. For sgRNA validation RPE WT or RPE p53^-/-^ BE3 expressing cells were infected with LentiGuide-puro-mCherry-NLS-*AAVS1* or LentiGuide-puro-GFP-NLS-sgRNA. For U2OS cells, 2 mg/mL puromycin was added 24 hours post infection and 48 hours after selection cells were plated for the experiment. For RPE WT and RPE p53^-/-^ BE3 cells 20 mg/µl puromycin was added 24 hours post infection and cells were plated 48 hours later for the experiment. GFP and mCherry expressing cells were plated 1:1 ratio on clear bottom 96 well plates at a density of 4,000 cells per well (2,000 per GFP or mCherry). 24 hours later cells were placed in media with or without drugs as in the base editing screen protocol and this was considered day 0. Drug or media was replaced every 3 days and cells were imaged prior to the media replacement. Cells were imaged on day 0, 3, 6, 9, 12, and 15. Plates were imaged on a Molecular Devices ImageXpress system, and the cells were scored as red or green using the MetaXpress imaging software.

### Viability assays

Short-term viability assays were completed with AlamarBlue (Invitrogen). HU, CPT or cisplatin was left in the media for 24 hours before being replaced with fresh media. AlamarBlue Cell Viability Reagent was added 72 hours after treatment and readings were taken at 590 nm 1-4 hours later. All viability measurements are presented as percent of the untreated control. Each assay was completed in triplicate. Cells stably expressing the RADX mutants of interest were plated for long term clonogenic survival. Colonies were allowed to grow for approximately two weeks prior to scoring with methylene blue staining (48% methanol, 2% methylene blue, 50% water). All clonogenic survival assays were completed in triplicate.

### Protein purification

His-MBP-RADX, -RADXOB2m and -RADXQVPK were purified from baculovirus infected *Sf9* cells as described in [15]. For purification of other RADX mutants, HEK293T cells were transfected using FLAG-GFP-RADX, or FLAG-GFP-RADX mutant constructs. Cells were lysed in NETN buffer (20 mM Tris pH 8.0, 150 mM NaCl, 1 mM EDTA, 0.5% NP40, cOmplete protease inhibitor cocktail) for 30 minutes at 4C. Clarified lysates were incubated with Anti-FLAG M2 magnetic beads (Sigma) for 2 hours at 4C. Beads were washed 2X in lysis buffer, 2X in LiCl buffer (lysis buffer containing 0.3 M LiCl) and twice in elution buffer (50 mM Tris pH 8.0 10% glycerol, 200 mM KCl, 1.0 mM EDTA, 1 mM DTT, 0.2 mM PMSF). The bound proteins were eluted in the elution buffer containing 0.25 mg/mL Flag peptide on ice for 90 minutes. RAD51 was purified as described previously [15].

### Immunofluorescence

Cells were plated in 96-well clear-bottom plates. Cells were incubated with media containing 10 mM EdU for 20 minutes, pre-extracted for 5 minutes on ice in 20mM HEPES, pH 7.0, 50mM NaCl, 3mM MgCl_2_, 300mM sucrose, and 0.5%Triton X-100 followed by fixation in 3% paraformaldehyde. Cells were blocked for 1 hour in PBS containing 5% BSA. EdU was labeled by addition of 2 mg/mL sodium ascorbate, 2 mM copper sulfate, and 5 mM Alexa Fluor 647 conjugated azide in PBS for 30 minutes. Primary antibody incubation was performed for 1 hour in 1% BSA in PBS followed by a 45-minute incubation in secondary antibody. Nuclei were stained with a 5-minute incubation with DAPI in PBS. Plates were imaged on a Molecular Devices ImageXpress system and integrated nuclear intensity of γH2AX of EdU-positive cells quantitated using the Molecular Devices software.

### Proximity ligation assay

Proximity ligation assay using antibodies to RADX and an anti-biotin antibody to recognize EdU after conjugation to a biotin-azide were completed according to the manufacturer’s protocol (Sigma) and images obtained and quantified using a Molecular Devices ImageXpress instrument.

### DNA molecular combing

Cells were labeled with 20 μM CldU (Sigma, C6891) followed by 100 μM IdU (Sigma, l7125), for the time indicated. Cells were embedded in agarose plugs and the assay was performed as per Genomic Vision’s manufacturer instructions. The DNA was stained with antibodies that recognize IdU and CldU (Abcam Cat#ab6326, BD Cat#347580) for 1 h, washed in PBS, and probed with secondary antibodies for 45 min. Images were obtained using a 40X oil objective (Nikon Eclipse Ti). Analysis of fiber lengths performed using Nikon Elements software.

### Immunoblotting

Whole-cell lysates were extracted using Igepal lysis buffer (50 mM Tris pH 7.4, 150 mM NaCl, 1% Igepal CA-630, 1 mM EDTA, pH 8.5) enriched with sodium fluoride (1 mM), sodium vanadate (1 mM), protease inhibitor cocktail (Roche), 25 units of Pierce Universal Nuclease, and 1 mM MgCl_2_. Proteins were analyzed by SDS-PAGE and immunoblotting.

### RADX-RAD51 pulldown assays

For the RAD51 pulldown, RAD51 mouse monoclonal antibody (14B4, Abcam) was incubated with protein A magnetic beads for 1 hour at room temperature. The resin was washed 1X in RAD51 binding buffer (50 mM Tris pH 8.0, 200 mM NaCl, 2 mM CaCl_2_, 2 mM ATP, 1 mM DTT). 100 nM of purified RAD51 was added to the beads and incubated further for 1 hour at room temperature. The resin was washed 2X in binding buffers before the addition of the purified RADX protein for 1 hour. Resin was washed 2X in binding buffer before the addition of 2X SDS buffer and resolved by SDS PAGE and immunoblotting.

### DNA binding assays

Dynabeads T1 (Life Technologies) were washed twice in TE buffer and bound to biotinylated poly dT50olignonucleotide at room temperature for 20 minutes. Beads were washed twice again in TE buffer followed with two washes in binding buffer (50 mM Tris pH 7.5, 250 mM NaCl, 1 mM DTT, 10% glycerol). 1 nM of bound DNA was added to 50 nM of RADX proteins and incubated at room temperature for 20 minutes. The beads were boiled in 2X sample buffer for 5 minutes, and eluted proteins were analyzed by immunoblotting.

### Co-immunoprecipitation

HEK293T cells were co transfected with either HA-RADX or Flag-GFP-RADX-mutants with 2 μg total DNA. 72 hours post transfection cells were harvested and lysed in CHAPS lysis buffer (50 mM Tris pH 7.5, 150 mM NaCl, 10% glycerol, 0.7% CHAPS, 5 μg/ml aprotinin, 5 μg/ml leupeptin, 1 mM sodium orthovanadate, 10 mM β-glycerol phosphate, 1 mM NaF, and 1 mM DTT) for 30 min on ice and cleared by centrifugation. Supernatants were incubated with anti-HA agarose beads for 1 hour at 4°C. Beads were then washed twice in CHAPS lysis buffer before the addition of 2X SDS sample buffer. Immunoprecipitated proteins were separated by SDS-PAGE and detected by immunoblotting.

### ATPase assay

The ATPase activity of RAD51 was measured using a NADH-coupled microplate photometric assay as described previously [15]. Briefly, RAD51 (3 μM) was incubated with 10 μM (nucleotides) of poly(dT) ssDNA for 5 minutes at 30°C prior to the addition of RADX (0-100 nM) or indicated RADX mutants (0-100 nM). ATP (3 mM) was included in the initial reaction with RAD51 and ssDNA. The reaction was allowed to proceed for 1hr at 30°C with readings at 340 nm every 30 seconds. A control reaction in which all reaction components were added except for NADH was used to subtract the background absorbance at 340 nm.

### Screen controls and analysis

For analysis of the data generated by NGS, pair-end reads were joined using PEAR to create a single read from the two paired end files [44]. Cutadapt was used to trim the sgRNA flanking sequences at the 5’ and 3’ ends [45]. The number of reads per sgRNA sequence was computed using MAGeCK count command [46]. The MAGeCK count summary demonstrated that the percentage of mapped reads was above 80% percent for all samples. The Gini index for all samples was below 0.047, which is an appropriate cutoff. Comparisons of T0 to T18 was performed using MAGeCK robust rank aggregation (RRA) using normalized values as the input.

### Statistical methods

All statistical analyses are described in the figure legends and were completed with Prism. Investigators were blinded to sample identities and all experiments were completed at least three times unless otherwise indicated.

## Abbreviations

DSB: double-strand break;
ssDNA: single-stranded DNA;
HR: homologous recombination;
OB-fold: oligosaccharide/oligonucleotide binding domain;
BE3: cytosine deaminase base editor;
NGS: next generation sequencing;
UGI: uracil DNA glycosylase inhibitor;
LFC: log2 fold change;
CPT: camptothecin

