## Supplemental figure for "CRISPR-dependent base editing screens identify separation of function mutants of RADX with altered RAD51 regulatory activity"

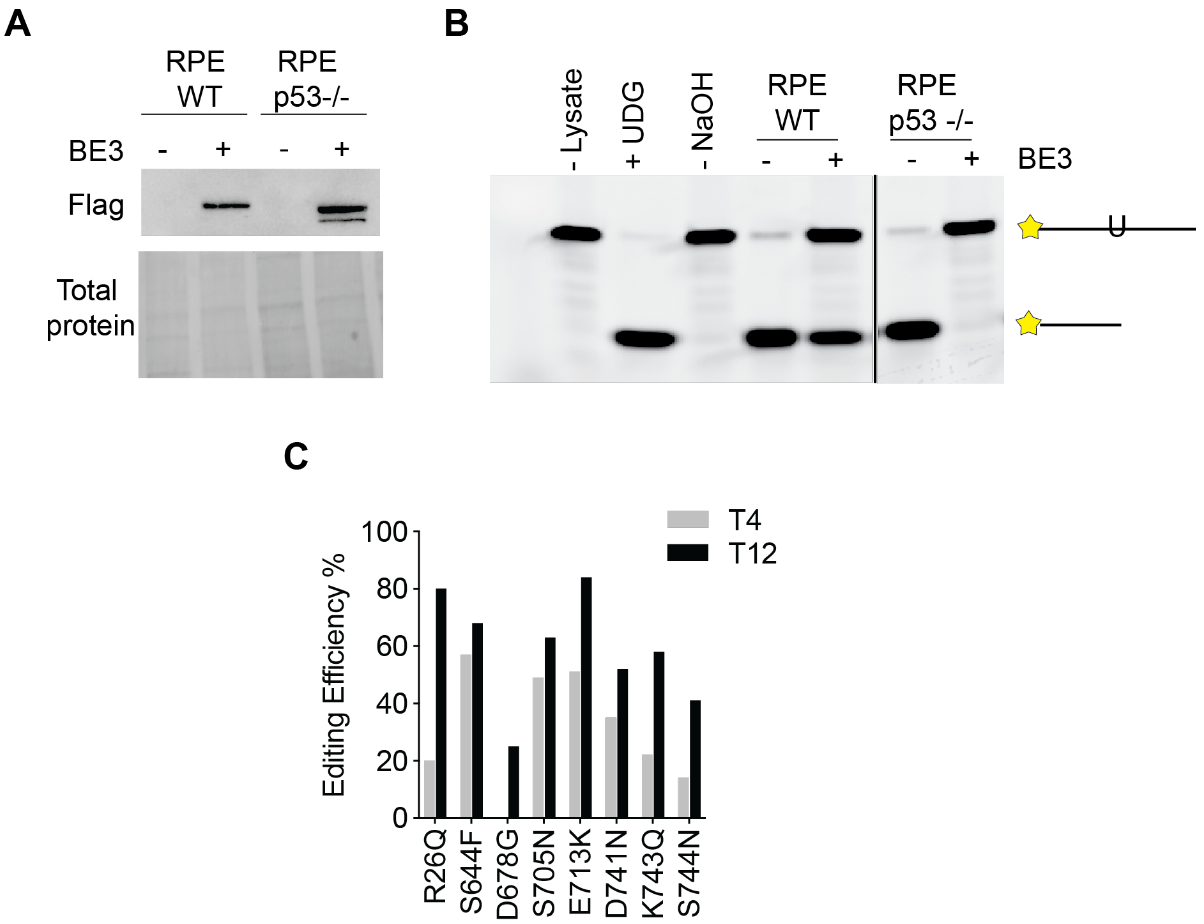


**Supplemental Figure 1. Quality controls for base editing screen. A)** Immunoblot showing Flag-BE3 expression in the RPE WT and RPE p53 -/- cells utilized for base editing screens **B)** Assay of UGI activity in BE3 expressing cells. **C)** Analysis of editing frequency induced by sgRNAs to indicated amino acid position at day 4 (T4) and day 12 (T12) post-selection as determined by Sanger sequencing and Inference of CRSIPR Edits (ICE) analyses [47].
